# Tuning metaplasticity in the adult visual cortex using flickering light

**DOI:** 10.64898/2026.02.16.706197

**Authors:** Francis Reilly-Andújar, Teresa L.M. Cramer, Eric Yuhsiang Wang, Nai-Wen Chang, Yi-Wen Chang, Arnold J. Heynen, Mark F. Bear

## Abstract

Synaptic connections in the brain are refined by sensory experience during an early postnatal critical period, but by adulthood synaptic connectivity is resistant to further changes. A consequence of lost plasticity is limited recovery from brain injury, disease, and adverse sensory experience. Thus, there is great interest in treatments that can promote synaptic modifications in the adult brain. In a wide variety of contexts, it has been established that the qualities of synaptic plasticity are not fixed but rather vary depending on the recent history of cellular or synaptic activity^1^. This plasticity of plasticity, or metaplasticity^2^ explains why temporary manipulations of brain activity (e.g., by drugs^3^, transcranial stimulation^4^, or sensory deprivation^5^) can set the stage for subsequent, potentially therapeutic, long-lasting synaptic modifications^6^. Here we tested the hypothesis that plasticity in the adult mouse visual cortex is influenced by prior exposure to temporally modulated light and discovered that different flicker frequencies have qualitatively different effects. Exposure to 60 Hz stimulation increased microglia density, depleted perineuronal nets (PNNs), and restored ocular dominance plasticity in response to brief monocular deprivation (MD). Exposure to 40 Hz flicker also enabled ocular dominance plasticity, but it did so in a distinct way and without PNN remodeling. A key distinction is that unlike 60 Hz flicker, which enabled depression of synaptic strength by MD, 40 Hz flicker promoted synaptic strengthening. Indeed, we found that 40 Hz flicker primed a rapid and robust recovery from the effects of long-term MD that failed to occur after 60 Hz flicker. Thus, metaplasticity can be non-invasively “tuned” by light flickering at different frequencies to encourage different forms of synaptic plasticity in the cerebral cortex, including modifications that enable recovery of function.

---

Our experiments were inspired by a previous discovery that repeatedly anesthetizing mice with ketamine restores ocular dominance plasticity in adulthood^7^. Obligatory for this effect are activation of cortical microglia and the digestion of PNNs that surround inhibitory neurons. Ketamine anesthesia is known to induce 40 Hz synchronous oscillations in the activity of cortical neurons (see, e.g., **Fig. S1**), and it has been reported that entrainment of activity with light at 40 Hz can also activate microglia in visual cortex^8^. These findings raised the intriguing possibility that exposure to 40 Hz flickering light might substitute for ketamine. Although 40 Hz flicker proved to be insufficient to cause degradation of PNNs, stimulation at a higher frequency—60 Hz—both activated microglia and depleted PNNs^7^. These surprising findings led us to hypothesize that 60 Hz flickering light (but not 40 Hz) can set the stage for subsequent experience-dependent plasticity. We note that this question is conceptually distinct from that addressed in a previous study showing that augmentation of visual experience with temporally coherent stimulation *during* a period of MD facilitates ocular dominance plasticity^9^.

We began by confirming that exposure to 40 Hz and 60 Hz flicker have distinct effects on microglia and PNNs. In these and all other experiments in this study, analyses were performed experimenter-blind to treatment condition. Adult (postnatal day (P) 74) C57BL/6J mice of both sexes received five sessions (one 2-hour session per day) of constant light (CL; no flicker), 40 Hz flicker, or 60 Hz flicker (**Fig. 1a)**. Microglial morphology in binocular primary visual cortex (V1b), assessed by ramification index and soma size, was altered after both 40 Hz and 60 Hz flicker compared to controls (**Fig. S2**). However, only 60 Hz flicker increased the overall number of microglia relative to CL mice (**Fig. 1b**), and elevated expression of Iba-1 and CD11b, an activation-associated microglial marker^10^ (**Fig. 1c; S2**). Aggrecan, a key component of PNNs that constrain ocular dominance plasticity^11^, was decreased by 60 Hz but not 40 Hz flicker (**Fig. 1d; Fig. S2**). Taken together, these experiments support the conclusion that while both 40 Hz and 60 Hz flicker elicit microglial changes, only 60 Hz flicker modulates microglia in a manner associated with PNN remodeling.

**Figure 1.**
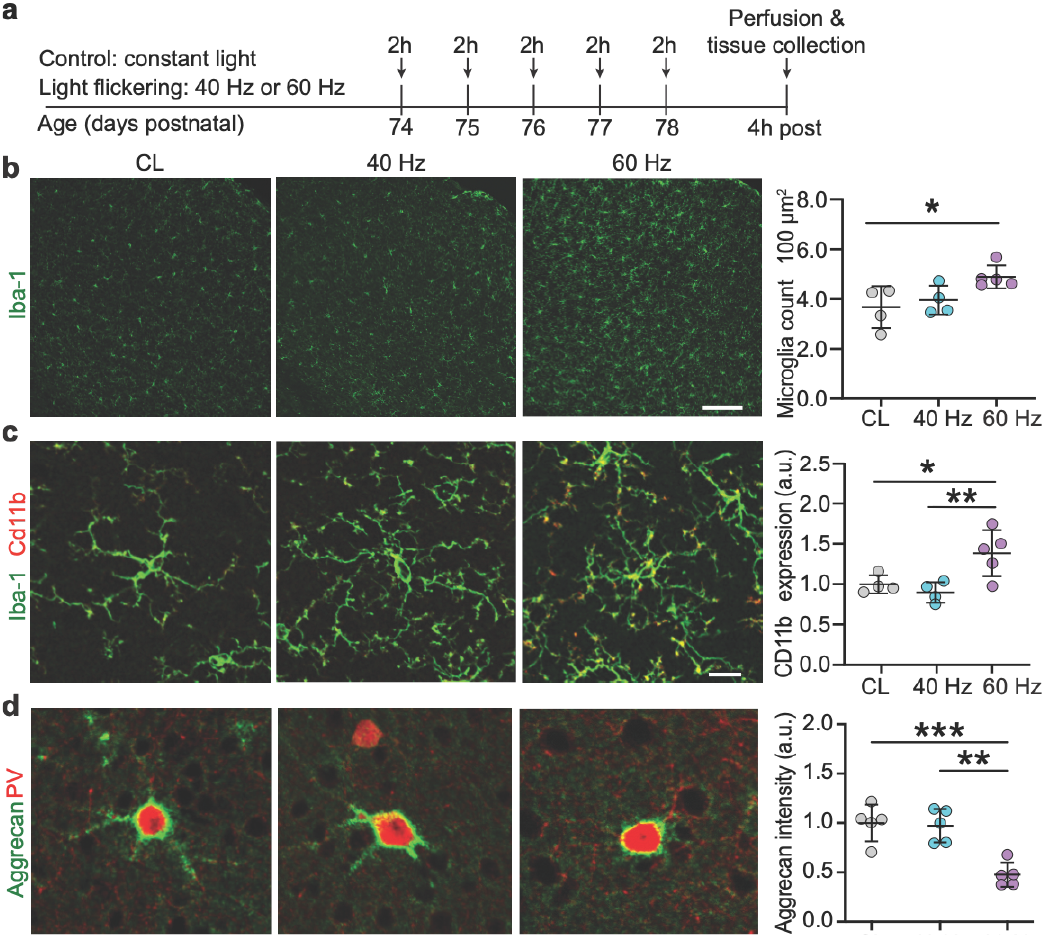
Light-flicker stimulation at different frequencies elicits distinct effects on microglia and PNNs in the adult primary visual cortex. **a**, Experimental timeline. **b**, Representative images and summary graph demonstrating increased counts of Iba1+ microglia in V1b of mice exposed to 60 Hz light-flicker compared to CL or 40 Hz light-flicker (n= 4-5 per group; CL: 3.673 ± 0.420; 40 Hz: 3.958 ± 0.289; 60 Hz: 4.892 ± 0.205; one-way ANOVA with Tukey’s test for multiple comparisons, p = 0.0365; CL vs 40 Hz: p = 0.8019; CL vs 60 Hz: p = 0.00395; 40 Hz vs 60 Hz: p = 0.1176). Scale bar 100µm. **c**, Representative images and summary graph demonstrating elevated CD11b expression in Iba1+ microglia in V1 of mice exposed to 60 Hz light-flicker as compared to CL or 40 Hz light-flicker stimulation (CL: 1.000 ± 0.056; 40 Hz: 0.900 ± 0.063; 60 Hz: 1.383 ± 0.127; one-way ANOVA with Tukey’s test for multiple comparisons, p = 0.0111; CL vs 40 Hz: p = 0.769; CL vs 60 Hz: p = 0.043; 40 Hz vs 60 Hz: p = 0.013). Scale bar 10µm. **d**, Representative images and summary graph demonstrating reduced aggrecan levels in V1b of mice exposed to 60 Hz light-flicker compared to CL or 40 Hz light-flicker stimulation (CL: 1.000 ± 0.083; 40 Hz: 0.9730 ± 0.076; 60 Hz: 0.477 ± 0.055; one-way ANOVA with Tukey’s test for multiple comparisons, p = 0.0003; CL vs 40 Hz: p = 0.9621; CL vs 60 Hz: p = 0.0007; 40 Hz vs 60 Hz: p = 0.001). Scale bar 20µm. Data are presented as mean ± SEM. *p < 0.05; **p < 0.01; ***p < 0.001.

Exposure to 60 Hz flickering light clearly leaves a trace in visual cortex, exemplified by the reduction in aggrecan-containing PNNs. A strong prediction from numerous lines of investigation is that degrading PNNs should restore a plastic response to brief MD that otherwise would be without effect after the critical period^7,12-19^. To test this hypothesis, adult (P78) mice received 4 days of MD after five daily sessions with flickering or constant light (**Fig 2a**). Ocular dominance in V1b was then assayed using single-unit recordings that enable measurements of the relative strength of inputs from the two eyes onto binocularly responsive neurons (**Fig. 2b**). For each unit, the maximal evoked responses to each eye were used to calculate an ocular dominance index (ODI; the contra-ipsi eye response difference divided by the sum), and the unit scores were averaged to yield an ODI score that represents the state of binocularity for that animal. Neurotypical mice have a positive ODI, reflecting the innate dominance of the contralateral eye. Depriving the contralateral eye of patterned vision during the critical period shifts the ODI to values closer to zero.

**Figure 2.**
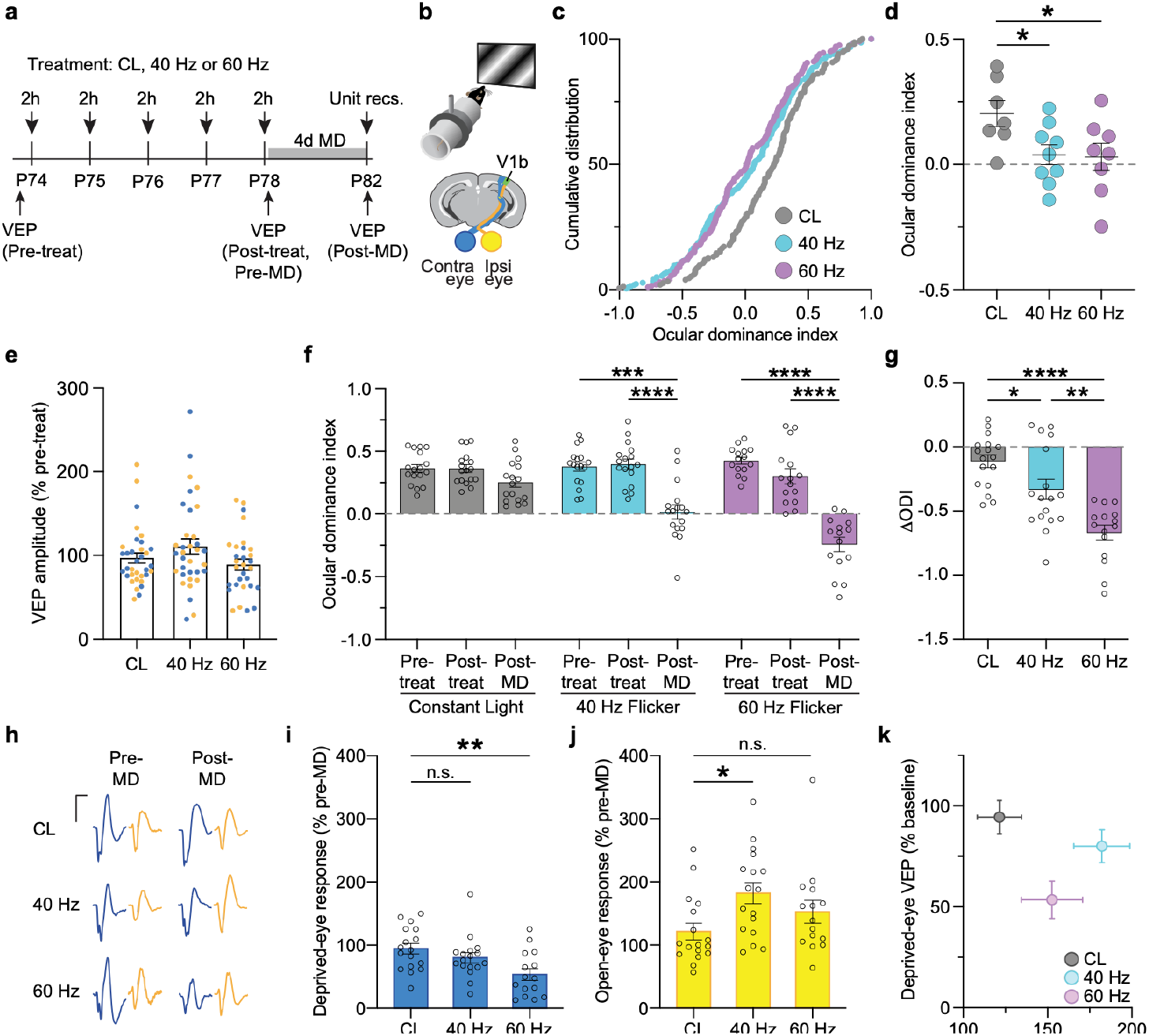
Exposure to 40 and 60 Hz light-flicker restores vulnerability to short-term monocular deprivation in adult mouse visual cortex through distinct mechanisms. **a**, Experimental timeline. In separate experiments, effects of MD were measured using acute single unit recordings with translaminar electrodes or VEPs from chronically implanted electrodes. **b**, Recordings were performed in V1b contralateral to the previously deprived eye (blue) in awake head-fixed mice viewing grating stimuli. **c**, Cumulative distribution of ODI values for single-units recorded across all layers of V1b in mice exposed to CL (n = 7), 40 Hz (n = 9), or 60 Hz (n = 8) flicker prior to 4 days of MD. Neurons from animals receiving either 40 Hz or 60 Hz light flicker show a statistically significant leftward shift in their ODI as compared to CL (control) animals. KS test with Bonferroni correction: CL vs 40 Hz: **p = 0.0014; CL vs 60 Hz: ****p < 0.0001; 40 Hz vs 60 Hz: p = 0.7592. **d**, Group comparisons (averaged unit ODI values per animal) indicate a statistically significant OD shift in animals receiving light flicker (CL: 0.2031 ± 0.053037, 40 Hz: 0.03926 ± 0.03970, 60 Hz: 0.02976 ± 0.05470; one-way ANOVA with Dunnett’s multiple comparisons test: p = 0.0386; CL vs 40 Hz: P = 0.0487, CL vs 60 Hz: p = 0.0420. **e**, Amplitudes of VEPs elicited through the ipsilateral (yellow) and contralateral (blue) eyes were not significantly altered by treatment with CL, 40 Hz, or 60 Hz flicker. **f**, ODI values from VEP recordings were unchanged by any treatment but shifted by MD in the flicker groups. Comparing ODI values across timepoints reveals no significant changes for the CL group (n = 17; Pre-Treat: 0.3634 ± 0.03115, Post-Treat: 0.3625 ± 0.03105, Post-MD: 0.2532 ± 0.03980, repeated-measures one-way ANOVA with Tukey’s multiple comparisons test, p = 0.0431; Pre-Treat vs Post-Treat: p = 0.9995, Pre-Treat vs Post-MD: p = 0.0949, Post-Treat vs Post-MD: p = 0.1270) whereas both 40 Hz and 60 Hz groups show a significant reduction in ODI values at the Post-MD timepoint (40 Hz, n = 17; Pre-Treat: 0.3769 ± 0.03648, Post-Treat: 0.3965 ± 0.04076, Post-MD: 0.01738 ± 0.05676, repeated-measures one-way ANOVA with Tukey’s multiple comparisons test, p < 0.0001; Pre-Treat vs Post-Treat: p = 0.7329, Pre-Treat vs Post-MD: p = 0.0002, Post-Treat vs Post-MD: p < 0.0001; 60 Hz, n = 15; Pre-Treat: 0.4247 ± 0.02797, Post-Treat: 0.3026 ± 0.05977, Post-MD: -0.2434 ± 0.05722, repeated-measures one-way ANOVA with Tukey’s multiple comparisons test, p < 0.0001; Pre-Treat vs Post-Treat: p = 0.1175, Pre-Treat vs Post-MD: p < 0.0001, Post-Treat vs Post-MD: p < 0.0001). **g**, Comparing the change in ODI from pre- to post-MD shows little effect of prior exposure to CL, but a significant effect of light-flicker. (CL: -0.1110 ± 0.04909, 40 Hz: -0.3302 ± 0.07826, 60 Hz: -0.6681 ± 0.05831, one-way ANOVA with Tukey’s multiple comparisons test, p < 0.0001; CL vs 40 Hz: p = 0.0428, CL vs 60 Hz: p < 0.0001, 40 Hz vs 60 Hz: p = 0.0016). **h**, Grouped average VEP waveforms for contralateral (blue) and ipsilateral (yellow) eyes across treatment condition and MD timepoint. Scale bar: 100 µV, 50 msec. **i**, Change in contralateral (deprived) eye VEP magnitude after MD (CL, 94.47 ± 8.293 %; 40 Hz, 80.02 ± 8.123 %; 60 Hz, 53.47 ± 9.258 %, one-way ANOVA with Tukey’s multiple comparison test, p = 0.0059; CL vs 40 Hz: p = 0.4466, CL vs 60 Hz: p = 0.0044, 40 Hz vs 60 Hz: p = 0.0864). **j**, Change in ipsilateral (non-deprived) eye VEP magnitude after MD (CL, 121.4 ± 13.13 %; 40 Hz, 182.2 ± 16.59 %; 60 Hz, 152.6 ± 18.16 %, one-way ANOVA with Tukey’s multiple comparison test, p = 0.0302; CL vs 40 Hz: p = 0.0227, CL vs 60 Hz: p = 0.3659, 40 Hz vs 60 Hz: p = 0.4055). Data are presented as mean ± SEM. *p < 0.05; **p < 0.01; ***p < 0.001. **k**, Summary of the different responses to 4d MD in the 3 groups.

As predicted, prior exposure to 60 Hz flicker enabled ocular dominance plasticity, as evidenced by a shift in ODI that was absent in adult mice exposed only to CL (**Fig. 2c-d**). Unexpectedly, however, we found that exposure to 40 Hz flicker also enabled the ocular dominance shift after MD. In both flicker groups, the shift in these adult mice was comparable in magnitude to what is observed in juvenile animals without flicker at the height of the critical period (**Fig. S3**).

Ocular dominance shifts after MD can be explained by depression of deprived-eye responses, potentiation of open-eye responses, or a combination of both. Deprived-eye depression occurs rapidly with ≤ 4 days of MD in 3-4-week old mice^20^, but fails to occur after ∼P32^21^. On the other hand, in both juvenile and young adult mice (up to 6 months of age^22^), periods of MD ≥ 7 days can trigger a potentiation of open-eye responses^20,23,24^. To gain insight into which of these mechanisms are facilitated by flicker exposure in adults, we monitored the absolute strength of V1b inputs from the two eyes using visual evoked potentials (VEPs) recorded from chronically implanted electrodes^25^. VEP recordings showed that although daily exposure to flicker alone produces no change in response magnitudes (**Fig. 2e**), subsequent 4-day MD shifts ocular dominance when preceded by exposure to *either* 40 or 60 Hz light flicker (**Fig. 2f-g**). However, analyses of the absolute VEP magnitudes indicate that the shift after 60 Hz flicker is caused primarily by depression of deprived-eye responses, whereas the shift caused by MD after 40 Hz flicker is mainly due to open-eye potentiation (**Fig. 2h-k**). Thus, exposure to one flicker frequency promotes *deprivation-induced* response depression while the other promotes *deprivation-enabled* response potentiation. By varying flicker frequency, we can tune subsequent plastic responses to the same environmental stimulus (in this case, MD).

One interpretation of flicker effects on ocular dominance plasticity is that 60 Hz stimulation uniquely releases a brake on youthful plasticity whereas 40 Hz merely accelerates the deprivation-enabled response potentiation that is already available in V1b of adult mice. Alternatively, it is possible that 40 Hz flicker additionally promotes forms of *experience-dependent* synaptic strengthening that are lost after the critical period. In species ranging from mice to human it has been shown that prolonged periods of MD initiated during the critical period cause the loss of visual responsiveness to deprived-eye input that does not recover spontaneously when binocular visual experience is restored in adults. We therefore asked if 60 or 40 Hz flicker exposure enable recovery from long-term (LT) MD.

Mice were monocularly deprived at the peak of the critical period (P28) for three weeks, given one week of normal binocular visual experience, and then randomly assigned to receive either no treatment, 40 Hz, or 60 Hz flicker each day for 5 days (**Fig. 3a**). Several days later, animals were prepared for single-unit recordings from V1b. As expected from previous studies^26,27^ in the absence of any treatment, restoring binocular visual experience alone failed to reverse the shift in ODI caused by long-term MD. To our surprise, there was also no indication that exposure to 60 Hz flicker enables recovery. However, mice in the 40 Hz flicker group showed a complete reversal of the ocular dominance shift, with ODI values comparable to those of normally reared mice of the same age (**Fig. 3b-c**).

**Figure 3.**
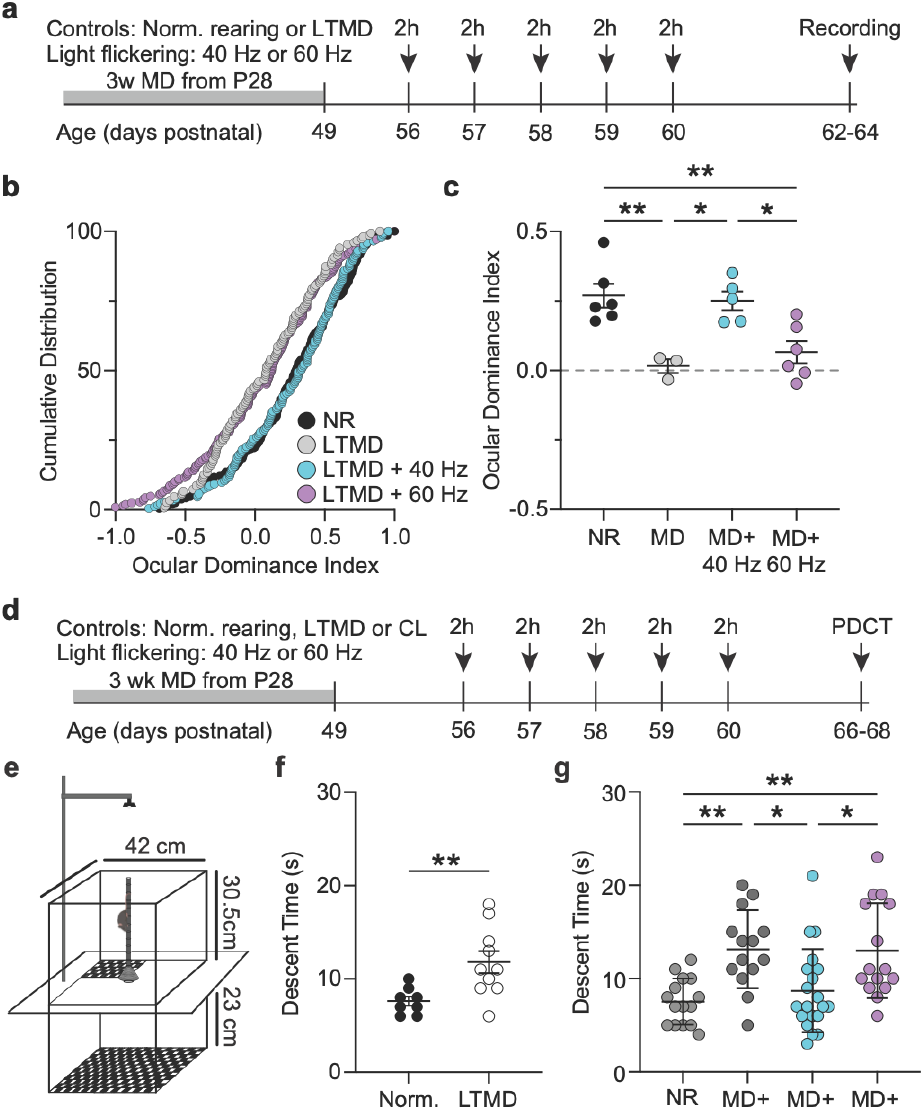
Electrophysiological and behavioral consequences of long-term MD are reversed after exposure to 40 Hz light-flicker. **a-c**, Single-unit recordings reveal that the ocular dominance shift caused by LTMD during the critical period is reversed by exposure to 40 Hz light-flicker. **a**, Experimental timeline. **b**, Cumulative distribution of ODI values for visually responsive single-units in V1b in mice reared normally (NR) or subjected to LTMD. LTMD animals were either untreated or exposed to either 40 Hz or 60 Hz light-flicker. The shift in ODI caused by LTMD is apparent in untreated and 60 Hz groups, but reversed in the 40 Hz group (KS – test with Bonferroni correction: NR vs LTMD: p < 0.001; NR vs LTMD + 40 Hz: p = 0.71688; NR vs LTMD + 60 HZ: p < 0.001; LTMD vs LTMD + 40 Hz: p < 0.001; LTMD vs LTMD + 60 Hz: p = 0.56635; 40 Hz vs LTMD + 60 Hz: p < 0.01). **c**, Group comparisons (averaged unit ODI values per animal) demonstrate that the OD shift produced by LTMD is completely reversed in animals receiving 40 Hz light-flicker treatment (NR: 0.2694 ± 0.04277; LTMD: 0.01600 ± 0.02450; LTMD + 40 Hz: 0.2509 ± 0.03403; LTMD + 60 Hz: 0.06587 ± 0.03974; One-way ANOVA with Tukey’s test for multiple comparisons, p < 0.001; NR vs LTMD: p = 0.0052; NR vs LTMD + 40 Hz: p = 0.9860; NR vs LTMD + 60 Hz: p = 0.0060; LTMD vs LTMD + 40 Hz: p = 0.0120; LTMD vs LTMD + 60 Hz: p = 0.8600; LTMD + 40 Hz vs LTMD + 60 Hz: p = 0.0172). **d-g**, The pole descent cliff task (PDCT) reveals visual deficits after LTMD that are reversed after receiving 40 Hz light-flicker. **d**, Experimental timeline for PDCT experiments. **e**, Experimental setup. Mice were positioned on top of a pole with head facing downward and the time taken to descend and dismount was recorded. **f**, LTMD animals have increased pole descent time when compared to NR mice (NR: 7.625 ± 0.498 seconds, LTMD: 11.8 ± 1.181 seconds, Mann-Whitney test: p = 0.00882). **g**, LTMD mice exposed to light-flicker at 40 Hz have descent times comparable to NR levels whereas the descent time of LTMD + 60 Hz is comparable to LTMD animals (NR: 7.533 ± 0.6389 sec, LTMD + CL: 13.14 ± 1.119 sec, LTMD + 40 Hz: 8.700 ± 0.9976 sec; LTMD + 60 Hz: 13.00 ± 1.317 sec, one-way ANOVA with Tukey’s test for multiple comparisons, p < 0.001; NR vs LTMD + CL: p = 0.0035, NR vs LTMD + 40 Hz: p = 0.8475, NR vs LTMD + 60 Hz: p = 0.0039, LTMD + CL vs LTMD + 40 Hz: p = 0.0179, LTMD + CL vs LTMD + 60 Hz: p = 0.9997, LTMD + 40 Hz vs LTMD + 60 Hz: p = 0.0199). *p < 0.05; **p < 0.01.

A functional consequence of long-term MD is deprivation amblyopia, a severe visual disorder in children^28^. A feature of amblyopia is impaired stereopsis, and in mice this can be studied using a recently developed pole descent cliff task^29,30^. In this assay, mice navigate down a vertical pole suspended over a glass plate divided into four quadrants, of which three appear as a visual cliff (**Fig. 3e**). A sensitive measure of imbalanced binocular vision is the time it takes to descend the pole. We recently reported that mice with a history of long-term MD show a marked slowing of descent compared to neurotypical controls^30^, and this finding was confirmed here (**Fig. 3f**). We therefore used this assay to test the hypothesis that vision can be improved in amblyopic mice by exposure to flickering light. In close correspondence with the electrophysiological findings, we observed that exposure of mice to light flickering at 40 Hz, but not 60 Hz, enabled functional recovery of binocular vision in animals that had been rendered amblyopic by long-term MD (**Fig. 3d-g**).

Each of the immunohistochemical, electrophysiological and behavioral experiments presented so far indicate that exposure to flickering light produces changes in V1b that enable forms of synaptic modification in adults, and that the precise nature of these changes differ qualitatively depending on stimulation frequency. To see if this conclusion could be corroborated with an orthogonal experimental approach, we used transcriptomics to ask if flicker at different frequencies leaves distinct molecular traces in neurons of V1b. Sections of visual cortex were prepared from mice after exposure to CL, 40 Hz or 60 Hz flicker and subjected to *in situ* multiplexed detection of RNA using the Xenium 5k platform^31^. Combining all samples, we identified a total of 282,364 cells. Then using the cell-by-gene matrix of segmented nuclei and uniform manifold approximation and projection (UMAP) for dimensionality reduction, we identified seven distinct cell types (excitatory and inhibitory neurons, astrocytes, microglia, oligodendrocytes, oligodendrocyte precursor cells, and endothelial cells, **Fig. S4**). Although multiple cell types displayed nominally significant differences in gene expression, we focused on glutamatergic neurons with changes that survived false discovery rate correction (FDR ≤ 0.05) in our sample. In these cells, 37 genes showed differential expression (≥ 0.5 log_2_ fold change) in the flicker mice compared to constant light control. Of these, 4 were unique to 40 Hz and 29 were unique to 60 Hz (**Fig. 4a**). Of the genes affected by 40 Hz flicker, 3 with well-known associations with synaptic plasticity (Arc, App, and CaMKIIa) showed increased expression compared to CL (**Fig. 4b-d)**. Conversely, most genes affected by 60 Hz flicker (31 of 33) showed significant decreases compared to CL (**Fig. 4c-d**). Interestingly, of the 4 genes that were affected by both 40 and 60 Hz flicker, one showed increased expression in both groups (*Arc*), two showed decreased expression in both groups (*Slc17a7* and *Atp2a2*) and one, *Camk2a*, showed an increase after 40 Hz and a decrease after 60 Hz (**Fig. 4d**). Although the biological significance of these specific changes requires further study, this experiment provides supportive evidence that exposure to flickering light leaves traces in the visual cortex, and these changes differ qualitatively depending on flicker frequency.

**Figure 4.**
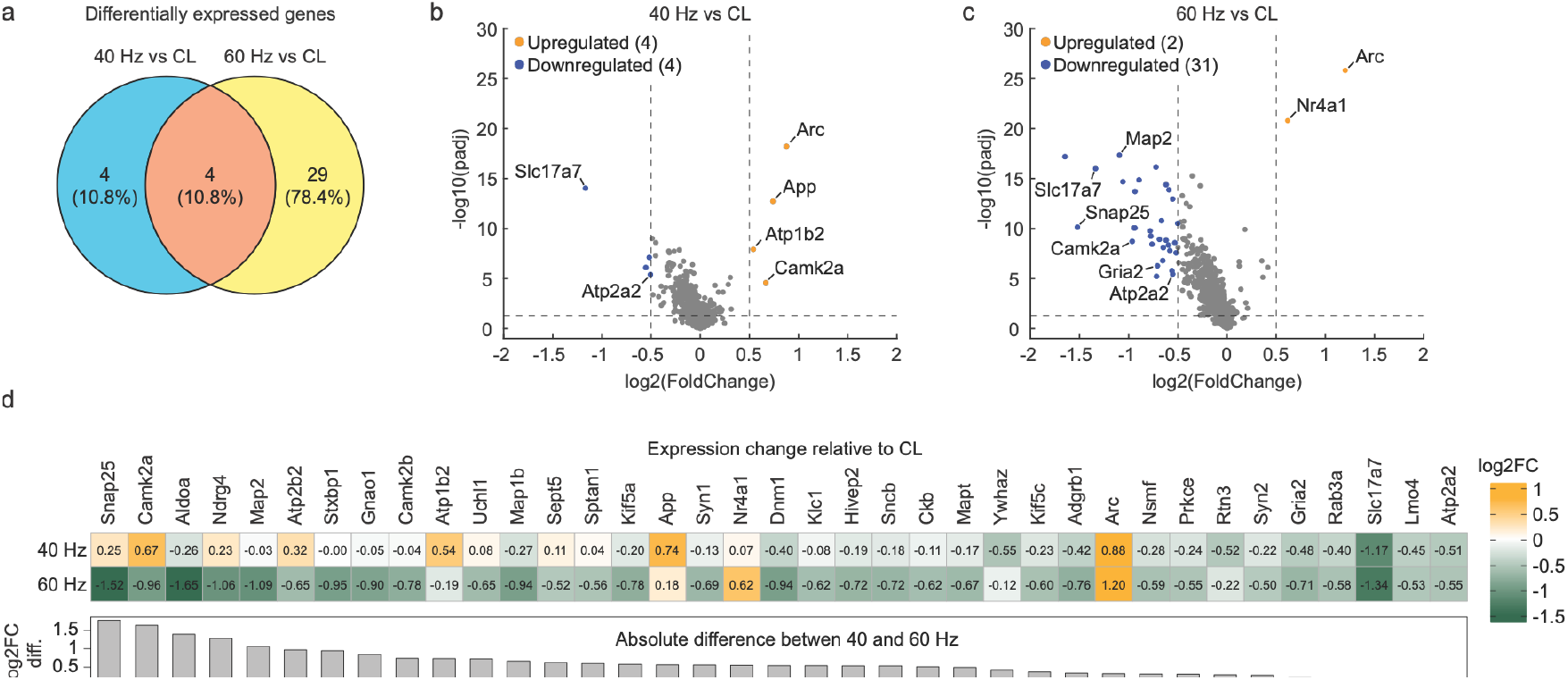
Qualitative differences in gene expression induced in glutamatergic neurons by light-flicker at 40 and 60 Hz. **a**, Venn diagram showing numbers of differentially expressed genes (DEGs) in glutamatergic V1b neurons in the flicker groups compared to CL. **b-c**, Volcano plots showing gene expression after treatment with 40 Hz or 60 Hz flicker compared to CL. Genes differing by ≥ 0.5 log2 fold-change and surviving a false discovery rate ≤ 0.05 appear in color. **d**, All genes significantly affected by flicker relative to CL in panels **b** and **c** are shown in descending order of the absolute difference in expression in neurons exposed to 40 and 60 Hz flicker. The heat map shows the difference in expression compared to CL.

Together, our data show that exposure of mice to light flickering at 40 or 60 Hz restores forms of synaptic modification in adult visual cortex that are normally restricted to an early postnatal critical period. While neither flicker frequency alone produces a lasting change in visual responses, both enable subsequent induction of modifications by manipulations of visual experience, thus satisfying the classical definition of metaplasticity^2^. However, the consequences of 40 and 60 Hz stimulation are quite distinct. Exposing mice to 60 Hz flicker renders connections from one eye vulnerable to synaptic depression in response to MD whereas 40 Hz stimulation enables the experience-dependent synaptic strengthening needed to reverse the effect of long-term MD.

While these experiments establish that flicker induces metaplasticity, *how* remains to be determined. In the case of 60 Hz flicker, a reasonable conjecture is that disassembly of PNNs enables changes in V1b inhibition^32,33^ that have been shown in young animals to allow the subsequent modification of excitatory thalamocortical synapses conveying information from the deprived eye^34,35^. This model is supported by abundant evidence that the late development of PNNs—specifically those surrounding PV+ interneurons—brings the critical period to a close^36,37^. We confirm here that microglia are increased in visual cortex of adult mice exposed to 60 Hz flicker which, in a previous study, was shown to be obligatory for the remodeling of PNNs^7,38^. Although there is an effect of 40 Hz flicker on microglia morphology, increased abundance and activation appear to be specific to 60 Hz. How microglial responses are tuned to different frequencies is unknown.

The mechanism behind the effect of 40 Hz flicker is also mysterious. The data do not support the hypothesis that remodeling of PNNs is the cause of enhanced synaptic strengthening, but this does not rule out lasting changes in inhibition. Indeed, in young adult mice it has been reported that “overstimulation” with 40 Hz light flicker (2x what was used here) for several days reduces the excitation-inhibition balance in V1^39^, but it is not clear how this change would promote subsequent experience-dependent strengthening of visual responses. Although the transcriptomic experiments show that 40 Hz stimulation uniquely triggers differential expression of some genes implicated in excitatory synaptic plasticity, the biological significance of these differences remains to be established. Additionally, our study was limited to the subset of genes available on the Xenium 5k platform and we are likely missing other genes of interest^40^. Nevertheless, a large and growing number of effects of 40 Hz sensory stimulation have been described since the seminal discovery that it can clear the brains of amyloid in mouse models of Alzheimer’s disease^41^. Recently it was discovered that 40 Hz flicker elevates adenosine^42^ and stimulates the expression and release of vasoactive intestinal polypeptide (VIP) from cortical interneurons which, in turn, induce glymphatic flow by enhancing arterial pulsatility^43^.

Interestingly, GABA release by VIP+ interneurons has been shown to be critical for the beneficial effects of locomotion on experience-dependent plasticity^44,45^. For any mechanism that is hypothesized, however, it must account for how treatment with 40 Hz flicker produces a persistent modification of the visual system that allows synaptic enhancement several days after the flicker has ceased.

Clearly additional research is needed to pin down how flicker at different frequencies modifies the cortex to promote synaptic plasticity. However, even without knowledge of the underlying mechanisms, the findings could have immediate practical impact. Visual stimulation at 40 and 60 Hz has been found to be safe and well tolerated by humans^46,47^. Thus, there is no obstacle to clinical testing of this approach to promote neurological recovery of function, most obviously in the context of vision loss caused by amblyopia.

## Acknowledgements

The authors wish to acknowledge the technical and administrative support of Hannah Manning and Nina Palisano. Research was supported by NIH (R01EY03773), the Freedom Together Foundation, the Severin Hacker Vision Fund, and the Swiss National Science Foundation. We thank the staff of the High-throughput Genomics and Big Data Analysis Core, Department of Medical Research, National Taiwan University Hospital for technical support, and acknowledge the spatial transcriptomic analyses provided by the National Genomics Center for Clinical and Biotechnological Applications of the Cancer and Immunology Research Center (National Yang Ming Chiao Tung University), the National Core Facility for Biopharmaceuticals (NCFB), and the National Science and Technology Council (NSTC).

## Methods

### Animals

All experiments were conducted on both female and male mice of the C57BL/6J background (The Jackson Laboratory). Mice were housed in groups of 2 – 5 same sex littermates after weaning at postnatal day 21 (P21). Mice were maintained on a 12h light-dark cycle (Light: 12 – 24 UTC; Dark 0 – 12 UTC) and had access to food and water *ad-libitum*. All procedures adhered to the guidelines of the National Institutes of Health and were approved by the Committee on Animal Care at the Massachusetts Institute of Technology.

### Light-flicker stimulation

LED light-flicker stimulation boxes were custom-built using black polycarbonate or ABS panels. Box dimensions were: 18 inches (length) x 15 inches (width) x 11 inches (height) and were specifically chosen to allow for a uniform distance of 4 inches from the LED strips to the walls of the mouse holding cage. An LED strip (RS Pro, 12V DC R,G,B White LED Strip Light, 4000k, Cat No. 265-1551) was wrapped on the inner sides of the stimulation box. LED light intensity was measured at the wall of the mouse cage at ∼850 Lux using a calibrated light meter (The Cooke Corporation, Cal-Light 400). LED light-flicker stimulation was delivered using an Arduino Uno device and consisted of a square-wave signal with a 50% duty cycle. Proper delivery of light-flicker stimulation was confirmed using an amplified photodetector (Thorlabs, PDA25K2). For all light-flicker stimulation experiments, mice (n = 2-5 per cage) were brought down to the behavioral space 30-60 minutes prior to the start of light-flicker stimulation. Light-flicker stimulation was initiated at 8 AM local time and finished at 10 AM. For experiments with a normally reared (NR; no light flicker) control group, mice were handled in the same manner as flicker conditions (i.e. 40 Hz, 60 Hz or Constant Light) with the exception that animals were not placed in a stimulation box.

### Chronic VEP recording implant surgery

Adult mice (postnatal day (P) 65 – 70) received a subcutaneous injection of buprenex hydrochloride (0.1 mg/kg) to provide pre-operative analgesia. Inhalation of isoflurane was used for induction (3% in O_2_) and maintenance (1-2% in O_2_) of anesthesia. Animals were maintained at 37°C for the duration of the surgical procedure. Prior to surgical incision the head was shaved and sterilized with rounds of povidine-iodide (10% w/v) and ethanol (70% w/v). The scalp was resected and the skull surface exposed. A steel headpost was placed on the skull surface, anterior to bregma, and secured using cyanoacrylate glue (Loctite 495 and 454). A small burr hole was made anterior to bregma and posterior to the headpost in order to place a silver-wire (A-M Systems) ground/reference electrode on the dural surface. Additional burr holes were made in both hemispheres at the sites of binocular visual cortex (∼3.0 - 3.1 mm lateral from the midline and slightly posterior to the lambda suture). A tapered 300 – 500 kΩ tungsten recording electrode (FHC) was lowered to a depth of ∼460 µm below the cortical surface, corresponding to layer 4, of each hemisphere. Electrodes were secured with cyanoacrylate glue. The remaining exposed skull surface was covered with a layer of cyanoacrylate glue and subsequently covered with dental cement (C&B-Metabond, Patterson Dental). Mice were monitored daily for the initial 72-hour post-operative period and given at least 5 days of recovery prior to the start of head-fixation habituation.

### Acute translaminar probe implant surgery

Surgical preparation including anesthesia/analgesia for acute translaminar probe recordings were performed as described for chronic VEP implants. The location of binocular visual cortex (∼3.1 mm lateral from the midline) was identified using stereotaxic procedures and the overlying skull demarcated for future craniotomy. A custom-built metal headplate was attached to the skull with cyanoacrylate glue to allow accessibility to binocular visual cortex on the day of acute recording. Dental cement (C&B-Metabond, Patterson Dental) was used to provide an additional level of coating on top of the cyanoacrylate glue. Exposed skull overlying V1b was covered using silicone adhesive (Kwik-Sil, World Precision Instruments). Mice were given 2 - 3 days of post-operative recovery prior to any form of head-fixation habituation. On the day of acute recording, under isoflurane anesthesia a craniotomy overlying V1b was performed using a small drill and consisted of thinning the skull until a thin layer could be removed with fine tip forceps. The exposed brain was subsequently covered with silicone adhesive and the mouse was given 1.0 – 1.5 hours of recovery prior to the start of recordings.

### Habituation to head-fixation

Mice were subjected to multiple 30-minute sessions of head-fixation habituation two days prior to the start of electrophysiological recordings. The habituation protocol consisted of placing the head-restrained mouse in the recording chamber and exposing it to an isoluminant gray screen, comparable to that of recording day. The initial sessions consisted of binocular viewing of the gray screen while the final sessions consisted of monocular viewing while habituating the mice to an occluder paddle.

### Monocular deprivation

Mice were anesthetized using inhalant isoflurane (3% in O2) and maintained with inhalant isoflurane (1 – 2% in O_2_). Medicated ophthalmic ointment (Neomycin, Polymyxin B and Bacitracin Zinc, Bausch + Lomb) was applied to the eyes to prevent dehydration during surgery and reduce the likelihood of infection following eyelid suture. The eye contralateral to the hemisphere of subsequent recording was sutured closed using 2 mattress stitches of sterile Prolene 7-0 (Ethicon). Sutures were tightened as necessary to ensure absence of light pinholes but not excessively in order to not alter the normal shape of the eye. After the MD period mice were anesthetized using inhalant isoflurane (3% in O_2_ induction,1-2% maintenance) and prior to eyelid re-opening a visual inspection was done to confirm proper eyelid closure. Sutures were removed and the eye flushed with drops of sterile saline (0.9% w/v) and inspection performed to confirm eye health. Mice were allowed to recover for at least 1 hour after eyelid reopening prior to the start of additional experiments.

### Visual stimulus delivery

Visual stimuli were created and delivered using custom written C++ and MATLAB scripts using the PsychToolbox extension (https://psychtoolbox.com). Chronic VEP recordings were obtained in response to each eye independently by covering the opposing eye with a light-tight occlude. To avoid potential order effects, we alternated the initial monocular viewing condition (contralateral or ipsilateral eye) per mouse. Mice were presented with 3 blocks of 50 phase reversals of oriented gratings at 0.2 cycles per degree, phase-reversing at 2 Hz and 100% contrast. Between every block of phase reversals there was a 30 second period of gray screen. Mice were presented with an orientation that differed by at least 30° from any previously presented orientation at each experimental timepoint to prevent confounding results due to the emergence of stimulus-selective response plasticity (SRP) ^48^. For acute translaminar probe recordings of single units, mice were exposed to 4 blocks of 50 phase reversals of oriented gratings at 0.2 cycles per degree, phase reversing at 2 Hz and 100% contrast viewed through each eye. There was a 30 second period of gray screen between each block of phase reversals. Oriented drifting gratings were also delivered at a spatial frequency of 0.2 cycles per degree and 100% contrast. The orientations for the drifting gratings were: 0°, 45°, 90°, 135°, 180°, 225°, 270° and 315°, each presented in randomized order for one second with a one second gray screen interleaved between each drifting grating. A total of 30 presentations were delivered for each oriented drifting grating.

### Electrophysiology

All electrophysiology was conducted in awake, head-fixed mice. The local field potential (LFP) recordings were sampled at 1 kHz, amplified and digitized using the Recorder-64 system (Plexon Inc.), a 300 Hz low-pass filter was applied. For ocular dominance plasticity experiments, VEPs were extracted and analyzed using the “VEPAnalysisSuite” developed by Jeffrey Gavornik (https://github.com/jeffgavornik/VEPAnalysisSuite'). Ocular dominance was expressed using the ocular dominance index = (Contra – Ipsi)/(Contra + Ipsi). Acute translaminar probe recordings were conducted using a 64-channel laminar probe (ASSY-77 H3, Cambridge Neurotech). The electrode was gently lowered (100 µm/min) into binocular primary visual cortex. To ensure proper targeting of binocular primary visual cortex we monitored visually evoked potentials elicited by stimulation of the ipsilateral eye by phase-reversing oriented gratings while the electrode traversed cortex. Recordings were amplified and digitized using the RHD Recording System (Intan Technologies). Recordings were sampled at 25 kHz and 0.1 Hz high-pass and 7.5 kHz low-pass filtered. Local field potential data was imported and analyzed using custom MATLAB scripts and the Chronux toolbox ^49^. Raw 25 kHz data from each channel was extracted and converted to µV. Data were downsampled to 1000 Hz and a third order 1 to 300 Hz Butterworth filter was applied. To account for any DC offset in the system, the mean of the entire channel’s data was subtracted from each time point. Data were locally detrended using the locdetrend function from the Chronux toolbox using a 0.5 sec window sliding in portions of 0.1 sec. Frequencies in the 58-62 Hz range were notched filtered using a third-order Butterworth. Custom MATLAB scripts based on the ‘spikes’ repository from the ‘cortex-lab’ (https://github.com/cortex-lab/spikes/blob/master/preprocessing/applyCARtoDat.m) were used to perform common median referencing of our LFP data. The output from the common median referencing was run through Kilosort 2.5. Units identified using Kilosort were curated using an interactive visualization tool (Phy, https://github.com/cortex-lab/phy). We used custom MATLAB scripts to extract data from our curated Kilosort output to perform unit analyses. We defined units as visually responsive if the averaged firing rate during drifting grating stimulation through either eye, elicited a response that was two times (2x) greater than the averaged firing rate observed in the 150 ms period prior to visual stimulation (150 ms pre-stim vs 150 ms post-stim). For units that met these criteria, we calculated their corresponding ocular dominance index values using the following equation: ODI = (Contra – Ipsi)/(Contra + Ipsi).

### Immunohistochemistry

Mice were anesthetized with Fatal-plus (50 mg/kg, i.p.) and transcardially perfused with oxygenated, ice-cold artificial cerebrospinal fluid (ACSF; 125 mM NaCl, 2.5 mM KCl, 1.25 mM NaH2PO4, 26 mM NaHCO3, 25 mM D-glucose, 2.5 mM CaCl2, 2 mM MgCl2). Following rapid brain removal, tissue was immersion-fixed in 4% paraformaldehyde for 90 min at 4 °C, rinsed in PBS, and cryoprotected overnight in 30% sucrose in PBS. Brains were sectioned at 40 μm on a sliding microtome and stored in cryoprotectant solution until use. Sections spanning V1 were processed for immunohistochemistry. After three 10-min washes in Tris-Triton buffer (50 mM Tris, 150 mM NaCl, 0.05% Triton X-100, pH 7.4), slices were incubated overnight at 4° C with primary antibodies diluted in Tris-Triton containing 0.4% Triton X-100 and 2% normal goat serum. Primary antibodies used were: mouse anti-PV (1:1000; Sigma; Cat# MAB1572), rabbit anti-PV (1:1000; GeneTex; Cat# GTX134110), rabbit anti-Iba-1 (1:500; Wako; Cat# 019-19741, Clone B56), rat anti-Cd11b (1:100; Thermo Fisher; Cat# 14-0112) and mouse anti-Aggrecan (1:2000; Merck; Cat# MAB1581). After three 10-min washes, sections were incubated for 30 min at room temperature with Alexa Fluor 488- and Cy3–conjugated secondary antibodies (1:500; Jackson ImmunoResearch) diluted in Tris-Triton containing 0.05% Triton X-100 and 2% normal goat serum. Sections were washed 3 times for 10 min, mounted onto gelatin-coated slides, and cover-slipped using DAKO fluorescence mounting medium

### Microscopy and image analysis

Stained brain sections were imaged using a confocal laser scanning microscope (Olympus FV1000; IX81 inverted stand) with 405 nm and 559 nm excitation. Images were compared to a mouse brain atlas to confirm proper targeting of V1b. For quantitative analysis, Z-stacks were acquired either with a 20x air objective (NA 0.75; six optical sections, 1 μm step size) or with a 40x air objective (NA 0.95; 10 optical sections, 0.5 μm step size). All imaging parameters were kept constant across experimental groups. To reduce variability, three sections encompassing V1 were collected per animal, with data obtained from a total of 4 mice (CL group) or 5 mice (40 Hz and 60 Hz groups). Measurements were averaged across sections for each mouse. Microglial ramification was quantified using an open-source Sholl Analysis plugin in Fiji/ImageJ^50,51^. All other quantifications were performed using a custom Python-based script in Fiji/ImageJ (https://github.com/dcolam/Cluster-Analysis-Plugin)^52^. Representative images were processed in ImageJ. Statistical analyses were carried out in Prism (GraphPad).

### Pole descent cliff task

All pole descent cliff task (PDCT) experiments were conducted in a quiet, temperature-controlled room during the 12-hour light cycle (Light: 12-24 UTC) as described ^30^. The apparatus used for testing consisted of an open field behavioral box (42 cm x 42 cm x 30.5 cm) with black surround walls, to prevent visual cues from the behavior room, placed above museum glass. A round wooden dowel (pole) 3.5 cm in diameter was screwed into a 75 mL Erlenmeyer glass flask, 9 cm at the bottom, for a total combined height of 52 cm. The pole and flask were covered with anti-slip silicone sleeves, texturized with a shelf and cabinet grip liner and spray painted white. The flask was ballasted with sand and a platform was glued to the center of the bottom of the flask, which created 1.5 cm separation between the bottom of the flask and the museum glass. An interior platform covering three of the four quadrants, covered with a black-and-white checkerboard pattern (2.5 cm × 2.5 cm) was positioned 23 cm below the museum glass. Another platform, covering one quadrant of the glass, also had a checkerboard pattern and was placed directly beneath the glass (see **Fig. 3e**). The pole was positioned in the center of the museum glass. A wide-angle USB camera with infrared capability (ELP) was kept 65 cm above the floor of the behavior apparatus and connected to a computer to record each session (resolution: 1080p, frames per second: 30). Before testing day, mice were individually handled for at least two days, with each session lasting five minutes. On the day of testing, whiskers were shaved, and the mice were brought to the behavioral testing room, where they remained in their home cage for at least 30 minutes to habituate to the testing space. Before the start of each trial, mice were allowed to freely explore the open-field behavioral box for five minutes, without the pole present and with the checkerboard pattern directly beneath the glass in all quadrants. At the end of the habituation period each mouse was allowed to descend the pole once. During test trials, three of the four quadrants were positioned 23 cm below the glass and the location of the “safe” quadrant (directly below the glass) was rotated every 1 – 2 mice to prevent side preferences due to external cues. Mice were placed on the top of the pole facing downward to encourage descent. Videos were acquired for all trials to allow analysis by an experimenter blind to treatment and rearing conditions. The chamber and pole were cleaned with peroxigard after each trial to eliminate odor cues. Male mice were tested first, followed by females. Recorded videos were used for manual scoring of safe vs. unsafe quadrant exits and for quantifying the time taken to descend and step off the pole.

### Statistics and blinding strategy

Experimenters were blind to treatment and data were de-identified prior to data analysis. Unless otherwise specified, data are presented as mean ± SEM. A one-way ANOVA with Tukey’s test for multiple comparisons was performed for data shown on Fig. 1b, Fig. 1c, Fig 1d, Fig. 2f, Fig. 2g, Fig.2i, Fig.2j, Fig. 3c, Fig. 3g, Fig. S2b, Fig. S2c, Fig. S2d, Fig. S2f, Fig. S2g, and Fig. S2h. A Kolmogorov-Smirnov (KS) test was conducted on the cumulative distribution data presented in Fig. 2c, Fig. 3b,and Fig. S3b. A one-way ANOVA with Dunnett’s test for multiple comparisons was performed for data presented in Fig. 2d. Mann-Whitney test was performed for data in Fig. 3f and Fig. S3f. An exact permutation test was performed for data in Fig. S1.

### In situ spatial transcriptomics profiling

Mice at P74 were exposed to 2-hour sessions of CL, 40 Hz, and 60 Hz flicker for 5 days, and euthanized with cervical dislocation 2 hours after the final flicker session. Brains were harvested and placed on ice-cold matrices. A 2-mm section from AP–2mm to–4mm from bregma was collected. All brain sections were immersed in 4% PFA for 20 hours at 4 °C before paraffin embedding. A total 6 brain sections (obtained from 2 animals per light-stimulus condition) were embedded and precisely aligned in the same cassette. H&E staining was performed to confirm that all samples were at the desired coordinates. Spatial transcriptomic profiling was conducted using the Xenium In Situ platform (10x Genomics). Tissue sections were hybridized with the Xenium Mouse 5K Pan Tissue & Pathways Panel (v1.0.0), comprising 5,006 RNA targets. Primary image processing, decodification, and signal assignment were performed via the Xenium Onboard Analysis pipeline (v3.3.0.1). To optimize cellular resolution, cell boundaries were delineated using a multi-modal segmentation strategy: (1) physical boundary stain-derived segmentation; (2) cytoplasmic interior stain-assisted nuclear expansion; and (3) nucleus-based isotropic expansion. Our workflow yielded a high-density dataset of 282,487 cells (mean density: 0.237 cells per 100 µm^2^), achieving a median detection of 411 genes and 568 transcripts per cell. Approximately 46.5% of all detected transcripts were successfully assigned to segmented cellular compartments.

### Xenium data processing and spatial subsetting

The raw Xenium output, including transcript matrices, spatial centroids, and segmentation polygons, was integrated into the Seurat (v5) environment using the ReadXenium framework. Stringent quality control (QC) was applied to ensure the fidelity of downstream inferences: genes detected in fewer than 3 cells and cells with fewer than 200 unique features were excluded. To minimize artifacts from cellular stress or debris, cells with a mitochondrial genomic contribution exceeding 10% were removed. For a high-resolution investigation of frequency-dependent responses, we focused on a manually curated anatomical region of interest, designated as V1b, which was defined by integrating spatial coordinates with initial unsupervised clustering results.

### Cell-type identification via consensus marker

To achieve a robust taxonomy of cell types within the V1b subset, we utilized a dual-algorithmic consensus approach for differential gene expression (DEG) analysis: (1) Non-parametric Marker Identification: We employed the Wilcoxon Rank Sum test (FindAllMarkers) with a minimum detection frequency (min.pct) of 0.1 and a log-fold change threshold of 0.5. (2) Linear Modeling: To account for variance stability, we concurrently applied the limma package to fit linear models across cellular populations. Statistical significance was strictly defined by a Bonferroni-adjusted P-value < 0.05 and an absolute log_2-fold change (|log_2FC|) > 0.5. Only DEGs identified by both frameworks were prioritized for defining distinct cellular identities, with intersections visualized using VennDetail (UpSet plots).

### Frequency-dependent transcriptional signature analysis

To identify specific molecular responses to different visual stimuli, we performed comparative DE analysis between 40 Hz and 60 Hz flicker conditions. DEGs in this context were determined using the Wilcoxon rank-sum test (P_adj < 0.05; |log_2FC| > 0.5). To systematically characterize the magnitude of frequency-specific sensitivity, we implemented a divergence-based ranking strategy for heatmap visualization (ComplexHeatmap). Genes were ordered along the horizontal axis based on the absolute divergence of their mean expression between the two conditions (|Value_40Hz - Value_60Hz|).

### Spatial statistics and computational environment

Functional enrichment of DEGs was performed to elucidate the biological pathways modulated by flicker frequency. Tissue-scale spatial distributions and cellular neighborhood architectures were analyzed to evaluate the systemic impact of light stimulation. All computational analyses were executed in the R environment (v4.3.0) utilizing Seurat (v5.0), ComplexHeatmap (v2.15), and the tidyverse (v2.0) suite^53^.

**Figure S1.**
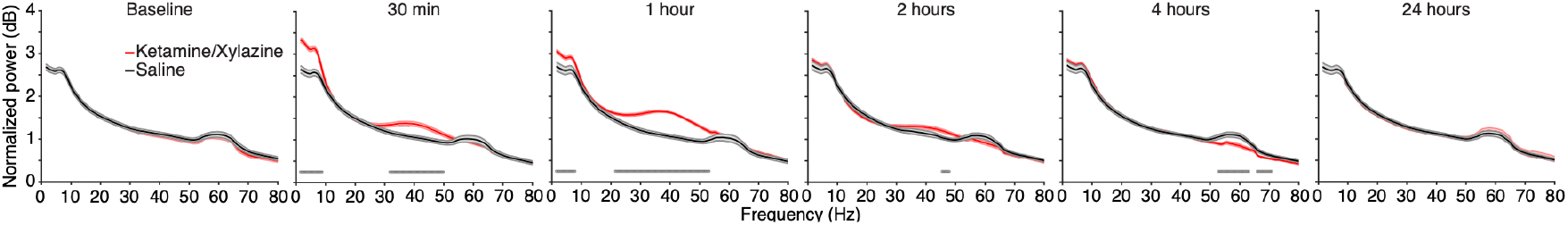
Ketamine anesthesia increases gamma power in the local field potential recorded in layer 4 of V1b. Mice were treated with ketamine/xylazine (100 mg/kg and 10 mg/kg i.p., respectively; n = 16) or saline (n = 11) and the local field potentials were recorded as animals viewed a gray screen. A statistically significant augmentation of gamma band (30 – 50 Hz) oscillations was observed from 30 min to 2 h following ketamine treatment compared to saline treated control animals. This same pattern of activation was observed across multiple, consecutive days of treatment (data not shown). Statistics were conducted via an exact permutation test. Significant differences (p val < 0.01) are indicated by marks near the x-axis. Shading represents mean ± SEM.

**Figure S2.**
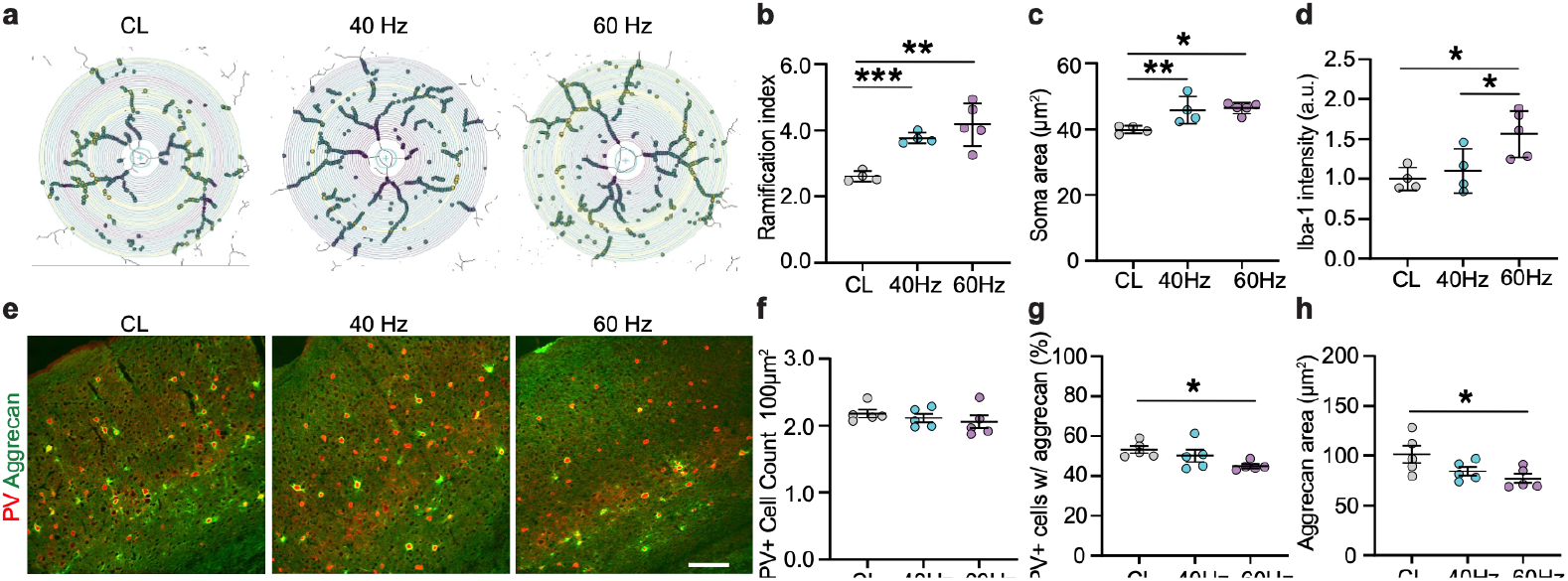
Light-flicker stimulation elicits differential effects on microglia and PNNs in the adult primary visual cortex. **a**, Representative Sholl overlay (concentric radii) on skeletonized microglia and summary graph (**b**) demonstrating increased ramification index in V1 of mice exposed to 60 and 40 Hz light-flicker compared to CL (n= 4-5 per group; CL: 2.661 ± 0.079; 40 Hz: 3.766 ± 0.082; 60 Hz: 4.175 ± 0.290; one-way ANOVA with Tukey’s test for multiple comparisons: p = 0.0009; CL vs 40 Hz: p = 0.008; CL vs 60 Hz: p = 0.00008; 40 Hz vs 60 Hz: p = 0.366). **c**, Summary graphs demonstrating increased soma size in V1 microglia of mice exposed to 60 and 40 Hz light-flicker compared to CL (CL: 39.84 ± 0.585; 40 Hz: 45.77 ± 2.082; 60 Hz: 46.41 ± 0.757; one-way ANOVA with Tukey’s test for multiple comparisons: p = 0.0079; CL vs 40 Hz: p = 0.023; CL vs 60 Hz: p = 0.009; 40 Hz vs 60 Hz: p = 0.928). **d**, Summary graphs demonstrating increased Iba-1 intensity in V1 microglia of mice exposed to 60 Hz light-flicker as compared to CL or 40 Hz light-flicker stimulation (CL: 1.000 ± 0.071; 40 Hz: 1.100 ± 0.139; 60 Hz: 1.563 ± 0.130; one-way ANOVA with Tukey’s test for multiple comparisons: p = 0.0149; CL vs 40 Hz: p = 0.840; CL vs 60 Hz: p = 0.0185; 40 Hz vs 60 Hz: p = 0.0494). **e**, Representative images and summary graphs (**f**) demonstrating unchanged PV+ cell counts across groups (n= 5 per group; CL: 2.180 ± 0.060; 40 Hz: 2.116 ± 0.063; 60 Hz: 2.059 ± 0.099; one-way ANOVA: p = 0.546). Scale bar 100µm. **g**, A reduced percentage of PV+ cells surrounded by aggrecan (CL: 53.18 ± 1.737; 40 Hz: 50.11 ± 3.104; 60 Hz: 43.45 ± 0.432; one-way ANOVA with Tukey’s test for multiple comparisons: p = 0.0175; CL vs 40 Hz: 0.463; CL vs 60 Hz: 0.016; 40 Hz vs 60 Hz: 0.098) and **h**, reduced aggrecan-covered area in V1 of mice exposed to 60 Hz light-flicker compared to CL or 40 Hz light-flicker stimulation (CL: 101.2 ± 8.637; 40 Hz: 84.41 ± 4.289; 60 Hz: 77.16 ± 4.650; one-way ANOVA with Tukey’s test for multiple comparisons: p = 0.0474; CL vs 40 Hz: 0.176; CL vs 60 Hz: 0.043; 40 Hz vs 60 Hz: 0.693). Data are presented as mean ± SEM. *p < 0.05; **p < 0.01; ***p < 0.001.

**Figure S3.**
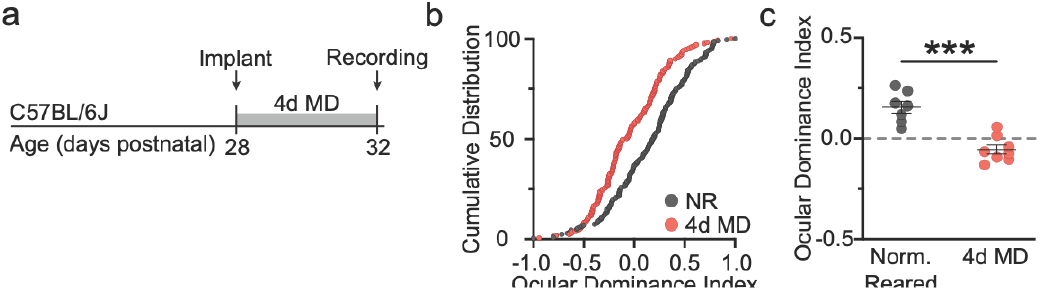
Ocular dominance shift caused by 4 days of MD during the critical period. **a**, Experimental timeline. **b**, Cumulative distribution of ODI values for all visually responsive (2x grey to preferred stimulus orientation) neurons across all layers of V1 of mice having normal binocular visual experience (NR) and mice having 4 days MD of the contralateral eye. Neurons from MD mice show a leftward shift in their ODI distribution signifying a shift in responsivity toward the fellow (nondeprived) eye. KS test: p < 0.001. **c**, Group comparisons (averaged single-unit ODI values per animal) show the expected OD shift in animals undergoing 4 days of MD (n = 8; -0.05459 +/-0.02216) as compared to control animals (n = 7; 0.1555 +/-0.02938; Mann-Whitney Test p = 0.0006). Data in **c** are presented as mean ± SEM. ***p < 0.001.

**Figure S4.**
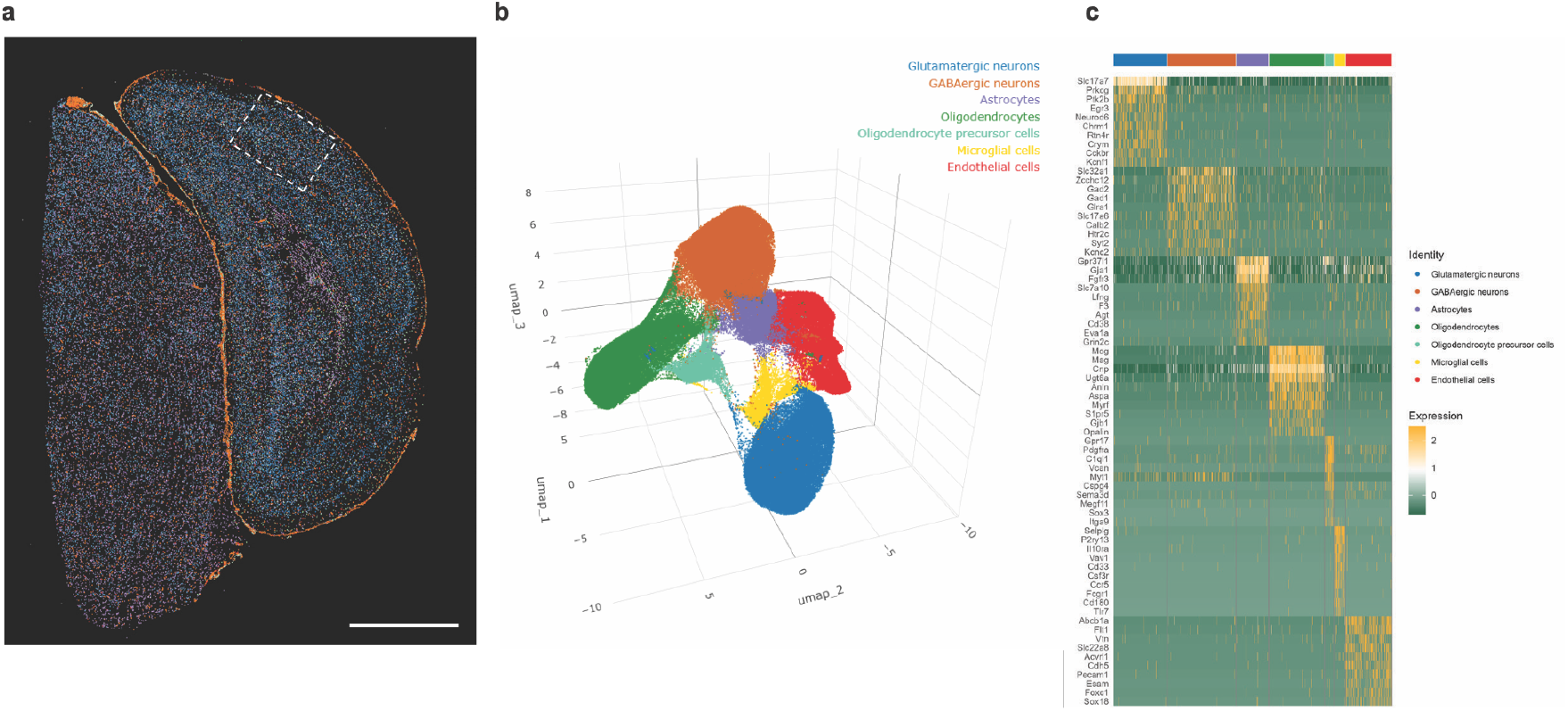
Transcriptomics. **a**, Image generated by 10x Genomics Xenium Explorer 3.2.0 of a section of mouse V1b subjected to analysis. Scale: 1 mm. **b**, Dimensional reduction analysis was performed on all defined cells and visually presented as UMAP. Seven cell types were defined based on differential expression of cell-type specific marker genes. **c**, Heatmap of cell-type specific markers for annotation. The top 10 genes enriched in each cell type are shown.

